# A specialized deceptive pollination system based on elaborate mushroom mimicry

**DOI:** 10.1101/819136

**Authors:** Satoshi Kakishima, Nobuko Tuno, Kentaro Hosaka, Tomoko Okamoto, Takuro Ito, Yudai Okuyama

**Affiliations:** Tsukuba Botanical Garden, National Museum of Nature and Science, Amakubo 4-1-1, Tsukuba, Ibaraki, 305-0005, Japan; Laboratory of Ecology, Graduate School of Natural Science and Technology, Kanazawa University, Kanazawa 920-1192, Japan; Laboratory of Insect Ecology, Faculty of Applied Biological Sciences, Gifu University, Yanagido 1-1, Gifu 501-1193, Japan

**Keywords:** *Arisaema*, deceptive pollination, Drosophilidae, floral scent, floral mimicry, mushroom

## Abstract

Despite its potential effectiveness for outcrossing, few examples of pollination via mushroom mimicry have been reported. This may be because the conditions under which the strategy can evolve are limited and/or because demonstrating it is challenging. *Arisaema* is a plant genus that has been suggested to adopt mushroom mimicry for pollination, although no compelling evidence for this has yet been demonstrated. Here, we report that *Arisaema sikokianum* utilizes mostly a single genus of obligate mycophagous flies (*Mycodrosophila*) as pollinators, and that the insect community dominated by *Mycodrosophila* is strikingly similar to those found on some species of wood-decaying fungi. Comparative chemical analyses of *Arisaema* spp. and various mushrooms further revealed that only *A. sikokianum* emits a set of volatile compounds shared with some mushroom species utilized by *Mycodrosophila*. Meanwhile, other closely related and often sympatric *Arisaema* species do not possess such typical traits of mushroom mimicry or attract *Mycodrosophila*, thereby likely achieving substantial reproductive isolation from *A. sikokianum*. Our finding indicates that mushroom mimicry is an exceptional and derived state in the genus *Arisaema*, thus providing an unprecedented opportunity to study the mechanisms underlying the coordinated acquisition of mimicry traits that occurred during a recent speciation event.

## Introduction

The diversity of floral forms found in angiosperms reflects the diverse plant–pollinator interactions that flowers engage in, wherein a particular suite of floral traits adapted to a specific functional group of pollinators is termed ‘pollination syndrome’ (Fenster et al. 2000, Ollerton et al. 2009). Among such relationships, a non-negligible proportion (associated with 7500 plant spp.) are based on deceptive pollination strategies (Renner 2006), in which plants lure pollinators by mimicking signals of food, mates or breeding sites without providing a reward to the pollinators. Although deceptive systems are generally considered ineffective at attracting pollinators compared with rewarding systems (Jersáková et al. 2006), the use of animal communities that are not otherwise involved in pollination may be advantageous because it allows escape from the ecological constraint of pollinator availability in time and space, as discussed in the context of the evolution of abiotic pollination (Cox 1991, Culley et al. 2002). Moreover, it may reduce the likelihood of heterospecific pollen transfer, as plants may trade pollinator visitation frequency for specificity (Scopece et al. 2010).

Mycophagous insects are rarely involved in pollination but are abundant in shady forest floor ecosystems, which in turn are relatively devoid of flower-visiting insects (Policha et al. 2016). Nevertheless, few examples of pollination via mushroom mimicry have been reported (Vogel 1978, Johnson and Schiestl 2016, Suetsugu 2018), and the well-established example of pollination by mycophagous insects via mushroom mimicry is limited to the orchid genus *Dracula*, in which mycophagous drosophilids are attracted to flowers that morphologically and chemically resemble mushrooms (Endara et al. 2010, Policha et al. 2016, 2019).

The rare occurrence of mushroom mimicry as a pollination system may indicate that the conditions under which the strategy can evolve are limited. For example, it may be because targeting specific groups of insects as pollinators is difficult, as most mycophagous insects have broad host ranges owing to the ephemeral nature of the fungi they feed on (Lacy 1984, Hanski 1989, but see Jonsell and Nordlander 2004). Moreover, demonstrating mushroom mimicry is often challenging, given that the floral traits associated with mushroom mimicry are not well understood, and the life histories of potentially mycophagous insects are largely unexplored.

In the present study, we explored an unverified example of specialized pollination via mushroom mimicry in an aroid species, *Arisaema sikokianum* Franch. et Sav., although its striking morphological similarities to mushrooms have long been recognized (see Figure 1*a*) (Vogel and Martens, 2000). *Arisaema* (Araceae) consists of 200 spp., with its center of diversity in E. Asia (Murata et al. 2018). The genus is well known for its ‘one-way traffic’ mechanism, with most members being sequential hermaphrodites (Vogel and Martens, 2000), in which the spathe functions as a trap for the pollinators (typically dipterans). Only male inflorescences have an exit hole at the bottom of the spathe, and pollinators are trapped in the spathes of the female inflorescence, which does not have an exit, after its pollination. Because mushroom-associated insects such as fungus gnats pollinate some *Arisaema* species, mushroom mimicry has been suggested as their attraction mechanism (Vogel, 1978; Vogel and Martens, 2000). However, such mimicry has not been demonstrated conclusively, and most species in the genus do not have morphological or chemical characteristics shared with mushrooms.

**Figure 1.**
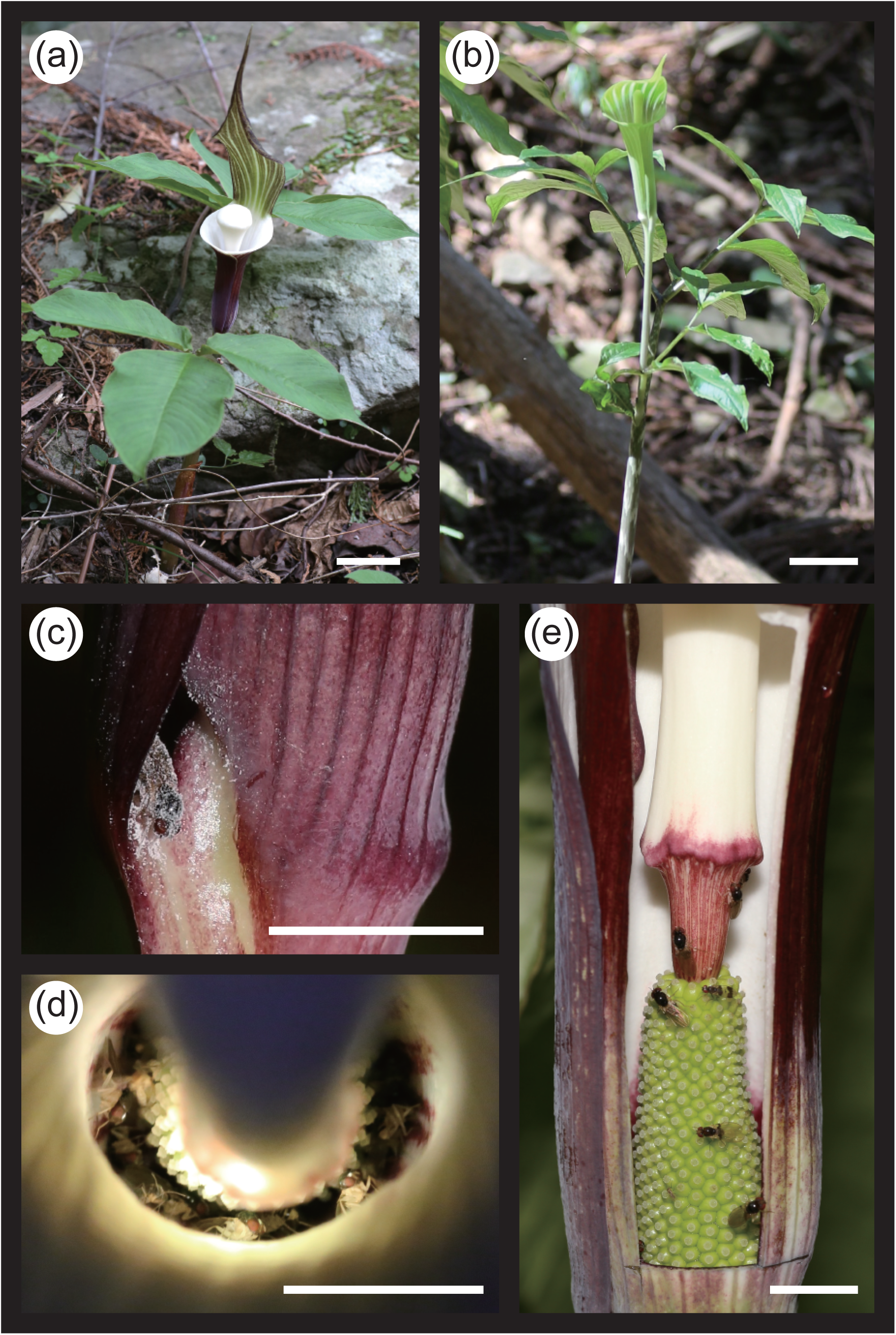
The flowering *Arisaema* spp. at the study sites. (a) *A. sikokianum*. (b) *A. japonicum*. (c) *Mycodrosophila* sp. escaping from the bottom exit hole of the male inflorescence of *A. sikokianum*. (d) Pollinators imprisoned in the female inflorescence of *A. sikokianum*, as visible from the top. (e) Pollinators imprisoned in the female inflorescence of *A. sikokianum*, as visible from the side after removal of the spathe. Scale = 5 cm (a and b) and 1 cm (c–e).

Here, we show that although not all *Arisaema* species have ‘mushroom flowers’, *A. sikokianum* has specialized floral traits for mushroom mimicry. To test the ‘mushroom flower’ hypothesis in *A. sikokianum*, we surveyed the pollinators trapped within the inflorescences and compared them with the insects attracted to various mushrooms. We further conducted comparative chemical analyses of floral and fungal volatile compounds to determine whether *A. sikokianum* imitates a specific fungus. Finally, we investigated whether the mushroom mimicry in *A. sikokianum* results in a unique pollination mode that enables reproductive isolation from other sympatric *Arisaema* species.

## Materials and Methods

### Target plants

*Arisaema sikokianum* is a tuberous perennial species endemic to western Japan and is ranked as vulnerable in the Red List of Japan (Ministry of the Environment, Government of Japan, 2012). This species is fairly rare across most of its distribution range and is ranked as highly endangered in most local (prefectural) red lists. However, the species is relatively abundant in Tokushima and Kochi prefectures. In mid to late April, the plant bears an inflorescence with a white, balloon-like appendix surrounded by a spathe that is white on the inside and dark purple on the outside (Figure 1*a*). As in most *Arisaema* spp., the inflorescence of *A. sikokianum* is unisexual, and the plant changes sex according to tuber size; larger tubers bear female inflorescences and smaller tubers male inflorescences (i.e., sequential hermaphroditism; Fukai, 2004). Male and female plants have almost identical inflorescences, except that a slit opening is present at the bottom of the inflorescence only in males, and the arrangement of the inflorescence tends to be higher in females. *Arisaema japonicum* Blume is closely related to *A. sikokianum* and belongs to the same section (*Pistillata*) (Ohi-Toma et al. 2016). The distribution ranges of *A. japonicum* and *A. sikokianum* overlap (Murata et al. 2018), and the two species sometimes exist sympatrically and bloom almost simultaneously. *Arisaema japonicum* has a similar growth habitat and ecology to *A. sikokianum* but is strikingly different in its floral characters, in that the inflorescence has a green, rod-shaped appendix surrounded by a green spathe (Figure 1*b*).

### Insect collection

We field-collected flower visitors trapped inside the spathe of the female inflorescence of >27 individuals of *A. sikokianum* from six locations during April 20–22, 2016 (sites 1–6; Figure 2; Table S1). Over the following years (2017–2018), we conducted additional field collections of flower visitors from the inflorescences of 6 female and 20 male individuals of *A. sikokianum* during April 23–25, 2017, and from the inflorescences of 4 female and 21 male individuals of *A. sikokianum* during April 18–20, 2018 at study sites 1 and 2 (Table S2). To collect flower visitors from the male inflorescences, the exit hole was plugged with cotton, and insect visits were allowed for 1–2 days. This experiment also enabled us to measure the daily flower visitation rates. Although it is difficult to discriminate the principal pollinators from other flower visitors in *Arisaema* spp., flower visitors that exit the male spathe are inevitably covered with pollen grains (Figure 1*c*) due to the structure of the inflorescence they must pass through. Accordingly, we identified the principal pollinators according to the following criteria: first, the insect species should visit both male and female inflorescences with substantial frequency; second, the insect should be small enough to pass through the exit hole of the male inflorescence; and finally, the insect should be large enough to carry a sufficient amount of pollen on its body.

**Figure. 2.**
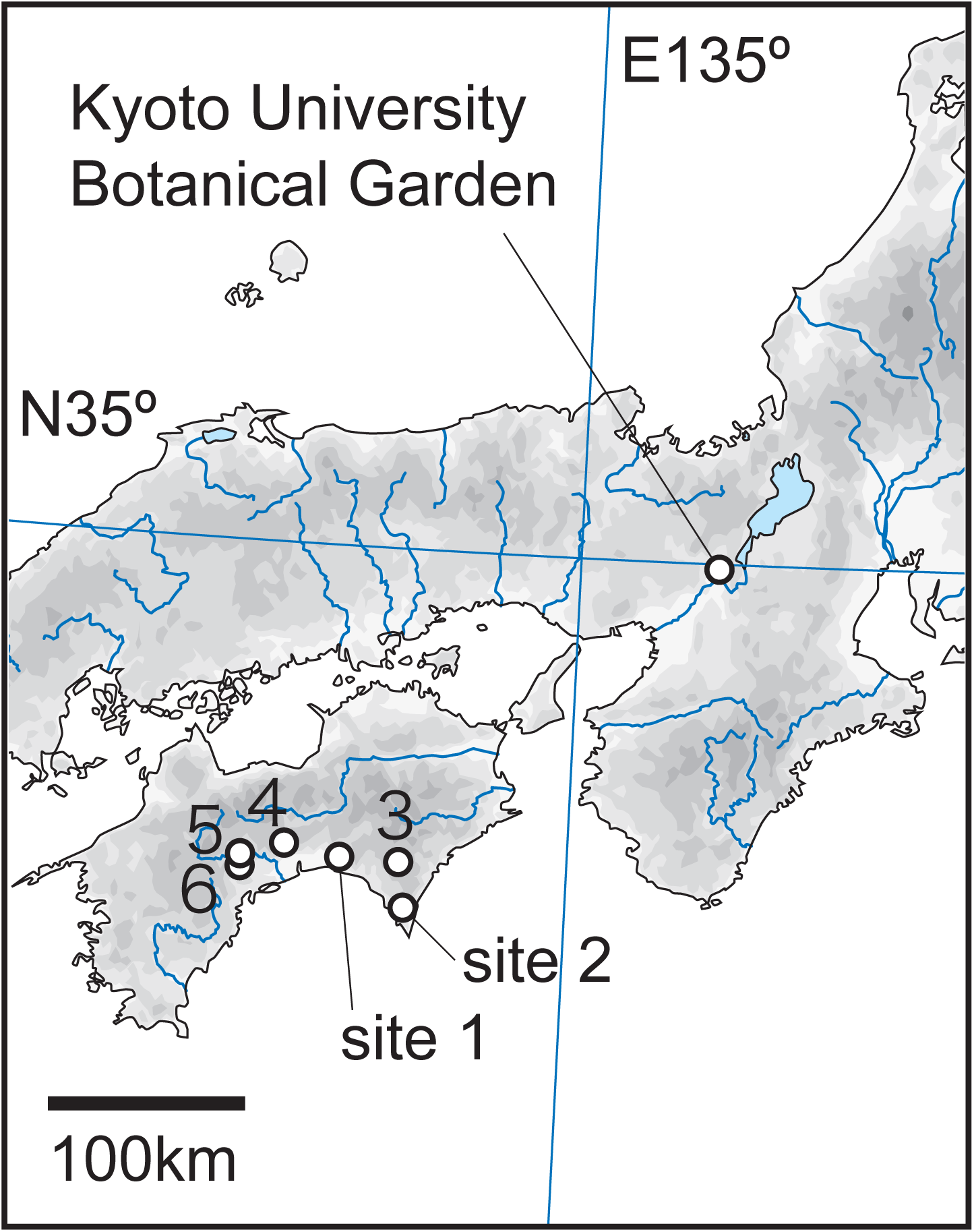
The location of the six field study sites and Kyoto University Botanical Garden.

To examine the role of the mushroom-like appendix of *A. sikokianum* in attracting pollinators, the swollen top half of the appendix was removed using scissors for 15 male individuals on April 23, 2017, and for 30 male individuals on April 11 and 12, 2019, at study sites 1 and 2. The bottom half of the appendix was left unremoved as it likely functions as a guard against insects escaping from the top of the spathe. The number of insect visits per day per inflorescence during the study period was compared between male individuals with a removed appendix (N = 45) and those with an intact appendix (N = 42).

To assess the presence of pollinator isolation, we also collected flower visitors from *A. japonicum* co-flowering with *A. sikokianum* at the same study sites. For *A. japonicum*, we collected insects from the inflorescences of 7 female individuals in 2016, 16 female individuals and 37 male individuals in 2017, and 15 female individuals and 34 male individuals in 2018 at sites 1 and 2 (Table S2).

Insects collected from the *Arisaema* plants were identified both morphologically and by DNA sequencing of a 710-bp fragment of the mitochondrial cytochrome c oxidase subunit I gene using a method described previously (Okuyama et al. 2018). All newly generated DNA sequences are deposited in the DNA Data Bank of Japan (DDBJ) under accession nos. LC193971–194126.

One of the authors (NT) extensively collected insects visiting the mushrooms at the Kyoto University Botanical Garden (Figure 2) and has reported the visitation pattern previously (Tuno, 2001). Thus, we reanalyzed the original data from that report (Tuno, 2001) to compare the insect communities on mushrooms and *Arisaema* inflorescences. In Tuno (2001), only mushroom-visiting dipterans were included in the analysis, and the species groups (an informal taxonomic rank narrower than a subgenus that is traditionally used in drosophilid taxonomy; O’Grady and DeSalle, 2018) rather than the individual species were used as the categories for analyses. We followed the same categorization scheme, but coleopterans and other insects were also included in the analysis. Coleopterans and other insects were each grouped into a single category because they represented only a minor portion (<10%) of the visitors to *Arisaema* inflorescences (see Results).

### Chemical analysis

We collected volatiles from *Arisaema* inflorescences or mushrooms using solid phase microextraction (SPME) headspace sampling with 50/30 µm divinylbenzene/carboxene/polydimethylsiloxane fibers (Supelco/Sigma-Aldrich, Bellefonte, PA, USA). Data on volatile composition obtained using SPME under standardized sampling conditions are highly repeatable, and comparable with absorbent-based trapping methods not only for assessing quality but also for quantity (e.g., Povolo and Contarini 2003, Friberg et al. 2013). Indeed, the volatile profiles of multiple individuals of *A. sikokianum* and *A. japonicum* sampled across different years were consistent (see Results), indicating that our results regarding the chemical similarity of volatiles emitted from *A. sikokianum* inflorescences and mushrooms is unlikely to be affected by the sampling method used; i.e., SPME vs. absorbent-based.

For *Arisaema* spp., the plants used for volatile sampling were collected from their native populations (nine *A. sikokianum* plants from four localities and eight *A. japonicum* plants from four localities, and one to four plants each of other *Arisaema* species) and were cultivated in the nursery in Tsukuba Botanical Garden for 0.5–3 years until the volatile sampling (Table S3). Fully open inflorescences were cut from a potted individual and immediately enclosed in a 500-mL or 1-L glass beaker capped with aluminium foil, and the SPME fiber was exposed to the headspace for 30 min. To clarify which parts of the inflorescences were responsible for the emission of volatiles in *A. sikokianum*, volatiles from the spathe, appendix and flower cluster were sampled separately by dissecting the inflorescence of a single *A. sikokianum* plant. For mushrooms, 30 fruiting bodies from 27 basidiomycous fungal species were collected, brought to the laboratory in Tsukuba Botanical Garden, and kept for < 2 days until the analysis (Table S4). Although these fungal species were sampled from the distribution area of *A. sikokianum*, they were selected because they are common species in the secondary deciduous forests of Japan where *A. sikokianum* and other *Arisaema* species grow. We aimed to sample a similar list of species to those studied in Tuno (2001), although our sample set was not identical due to the difficulty of comprehensively collecting these fungal species as well as uncertainty in the identification of some fungal species in Tuno (2001). The peak timing of fruiting body production varies among the fungal species sampled, but the majority are capable of producing fruiting bodies during the flowering season of *Arisaema* spp. (April–May, see Table S4). Each of the collected fruiting bodies was placed in a glass container of the appropriate volume (a 100-mL vial or a 500-mL or 1-L beaker). The ambient air of the sampling room was also analyzed several times during the sampling period to distinguish the volatiles emitted from the *Arisaema* plants or mushrooms. The sampled SPME fiber was analyzed immediately by gas chromatography–mass spectrometry. We used a GCMS-QP2010SE system (Shimadzu, Kyoto, Japan) equipped with an Rtx-5SilMS capillary column (30 m × 0.25 mm; film thickness, 250 nm; Restek, Bellefonte, PA, USA). Helium was used as the carrier gas at a velocity of 48.1 cm/s, and the injector temperature was 250°C. The injector was operated in the splitless mode for 1 min. Electron ionization mass spectra were obtained at a source temperature of 200°C. The oven temperature was programmed to the following sequence: 40°C for 5 min, followed by an increase to 220°C at 5°C/min and then 280°C at 15°C/min, at which the oven was held for 5 min.

For every volatile compound, the retention index was calculated using n-alkane (C6–C20) standards. Then, identification was carried out by comparison with the mass spectra in the NIST14 and NIST14s libraries and retention indices reported in the NIST Chemistry WebBook (Linstrom and Mallard 2001). Where possible, we further verified the identity of these compounds by comparing the mass spectra and the retention indices of the corresponding commercially available authentic compounds (purchased from Wako Pure Chemicals [Tokyo, Japan], Tokyo Chemical Industries [Tokyo, Japan], or Supelco/Sigma-Aldrich [Bellefonte, PA, USA]).

### Data analysis

To characterize the volatile profiles of the individual samples, the peak area of the total ion chromatogram was used. Samples with no detectable volatile compounds were removed from the analysis. The composition of each compound within a sample was scored by the relative peak area (percentage) to calculate the Bray–Curtis dissimilarity among the samples, and then nonmetric multidimensional scaling was performed to compare the volatile profiles among samples using the R packages vegan (Oksanen et al. 2016) and MASS (Venables and Ripley 2002). The compositions of the insects collected from *Arisaema* inflorescences and mushrooms were also analyzed using similar procedures.

To evaluate the effect of removing the swollen top half of the *A. sikokianum* appendix on the insect visitation rate, we used a generalized linear mixed model (glmer in the R package lme4) (Bates et al. 2015) using a Poisson error distribution and log link function. We fitted “treatment” as a fixed term and “years”, “populations” and “the length of survey periods” as random terms in the generalized linear mixed model.

## Results

### *Insect assemblages attracted to* A. sikokianum, A. japonicum *and fungal fruiting bodies*

In the three consecutive study years (2016–2018), we collected and identified a total of 266 and 101 insect visitors from female and male inflorescences of *A. sikokianum*, respectively, and 83 and 68 insect visitors from female and male inflorescences of *A. japonicum*, respectively. On average, 1.42 and 1.14 insect individuals visited per one male inflorescence of *A. sikokianum* per day, whereas 0.30 and 0.72 insect individuals visited per one male inflorescence of *A. japonicum* per day during the study periods in 2017 and 2018, respectively (Table 1). The removal of the swollen top of the appendix from the male *A. sikokianum* inflorescence resulted in a significant decrease in insect visits (N = 87, Z = 2.885, df = 82, P = 0.004, two-tailed): 0.82 and 1.51 insect individuals visited each male inflorescence per day in the treatment and control groups, respectively (Table 2).

**Table 1.** Pollinator visitation frequency to male inflorescences of *A. sikokianum* and *A. japonicum* across study sites and years.

| Species | Year | Site | No. of<br>inflorescences | Pollinator visits | Visits per<br>inflorescence | Visits per<br>inflorescence<br>per day |
| --- | --- | --- | --- | --- | --- | --- |
| *A. sikokianum* | 2017 | 1 | 10 | 32 | 3.20 ± 3.26 | 1.60 |
|  |  | 2 | 15* | 39 | 2.60 ± 4.44 | 1.30 |
|  |  | total | 25 | 71 | 2.84 ± 3.94 | 1.42 |
|  | 2018 | 1 | 6 | 12 | 2.00 ± 1.67 | 1.00 |
|  |  | 2 | 15 | 36 | 2.40 ± 1.72 | 1.20 |
|  |  | total | 21 | 48 | 2.29 ± 1.68 | 1.14 |
| *A. japonicum* | 2017 | 1 | 22 | 15 | 0.68 ± 0.99 | 0.34 |
|  |  | 2 | 15 | 7 | 0.47 ± 0.83 | 0.23 |
|  |  | total | 37 | 22 | 0.59 ± 0.93 | 0.30 |
|  | 2018 | 1 | 26 | 46 | 1.77 ± 2.39 | 0.88 |
|  |  | 2 | 8 | 3 | 0.38 ± 0.52 | 0.19 |
|  |  | total | 34 | 49 | 1.44 ± 2.18 | 0.72 |

\*Only a subset of inflorescences was used for identification of pollinators.

**Table 2.** Effects of appendix removal on pollinator visit frequency to *A. sikokianum*.

| Treatment | Year | Site | Study period | No. of<br>inflorescences | Pollinator<br>visits | Visits per<br>inflorescence | Visits per<br>inflorescence<br>per day |
| --- | --- | --- | --- | --- | --- | --- | --- |
| Control | 2017 | 2 | 23–25 Apr | 15 | 39 | 2.60 ± 4.44 | 1.30 |
|  | 2019 | 1 | 11–13 Apr | 7 | 18 | 2.57 ± 2.23 | 1.29 |
|  |  | 2 | 12–13 Apr | 20 | 35 | 1.75 ± 1.71 | 1.75 |
|  |  | total |  | 42 | 92 | 2.19 ± 3.01 | 1.51 |
| Removal | 2017 | 2 | 23–25 Apr | 15 | 39 | 2.60 ± 2.56 | 1.30 |
|  | 2019 | 1 | 11–13 Apr | 9 | 9 | 1.00 ± 1.32 | 0.50 |
|  |  | 2 | 12–13 Apr | 21 | 13 | 0.62 ± 0.86 | 0.62 |
|  |  | total |  | 45 | 61 | 1.36 ± 1.88 | 0.82 |

The composition of insect visitors found in the inflorescences differed strikingly between *A. sikokianum* and *A. japonicum* (Figure 3*a*). In *A. sikokianum*, 64.4–93.4% of the insect visitors were drosophilid flies (Figure 1*c–e*, Table S5). In particular, *Mycodrosophila* spp. accounted for 66.7% of the total visitors, of which the most abundant species was *Mycodrosophila gratiosa*, accounting for 37.2% of the total visits. The insect visitors to the inflorescences of *A. sikokianum* with the appendix removed were also dominated by *Mycodrosophila*, which accounted for 76.3% of the total visits. Meanwhile, in *A. japonicum*, Sciaridae (26.0%), Cecidomyidae (20.0%), and Mycetophilidae (14.7%) were the main insect visitors, while only one drosophilid individual, which was a species that has never been collected previously from the *A. sikokianum* spathe, was collected (Table S5). Accordingly, there was little overlap in pollinator assemblages between *A. sikokianum* and *A. japonicum*.

**Figure 3.**
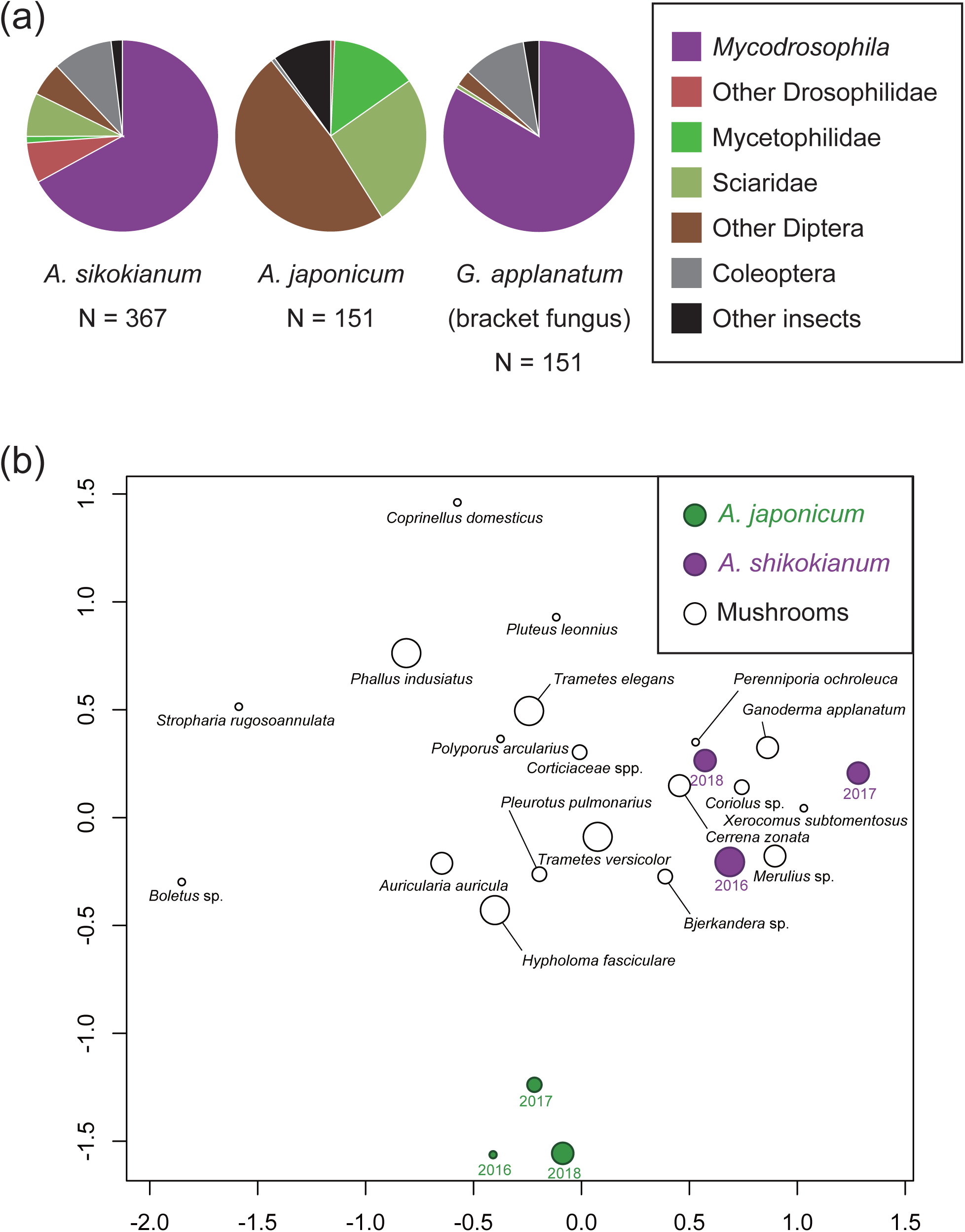
The insect assemblages that visited the inflorescences of *Arisaema* spp. and the various mushrooms. (a) Categories of insect species trapped inside the spathes of *A. sikokianum* and *A. japonicum* and those visiting the fruiting bodies of the bracket fungus *Ganorderma applanatum* illustrated as pie charts. (b) Nonmetric multidimensional ordination plot of the insect assemblages. The size of the plots corresponds to the sampling size (four levels: N < 40, N < 80, N < 160, and N > 160, respectively). The individual plots for *Arisaema* spp. correspond to the study years (2016–2018).

The insect assemblages trapped inside the inflorescences of *A. sikokianum* were strikingly similar to those attracted to the fruiting bodies of mushrooms, especially those of wood-decaying polyporous fungi (Figure 3*b*). We found that the level of dissimilarity of the compositions of attracted insects was much lower between *A. sikokianum* inflorescences and mushrooms (0.13–1.00, 0.61 on average) compared with between *A. japonicum* and mushrooms (0.61–1.00, 0.89 on average) or between *A. sikokianum* and *A. japonicum* (0.77–0.99, 0.88 on average).

### *Scent profiles of the inflorescences of* Arisaema *species and fungal fruiting bodies*

We detected a total of 94 volatile compounds (24–61 compounds per individual sample) in the headspace samples of nine *A. sikokianum* individuals (Table S6). Fifteen compounds were common among the samples. The common volatiles consisted of monoterpenes, sesquiterpenes, benzyl nitrile, 2-ethylhexyl acetate and C8 aliphatics. Benzyl nitrile and one of the C8 aliphatics, 3-octanone, were the most abundant, comprising 0.2–39.4% and 12.9–40.7% of the total peak area of the volatiles detected, respectively (Figure 4*a*; Tables 3 and S6). *Arisaema sikokianum* has a showy white appendix that resembles a fungal fruiting body (mushroom). We confirmed that the mushroom-like appendix emits a major portion of the volatiles detected from the inflorescence, although the spathe had a similar volatile profile (Figure S1).

**Table 3.**
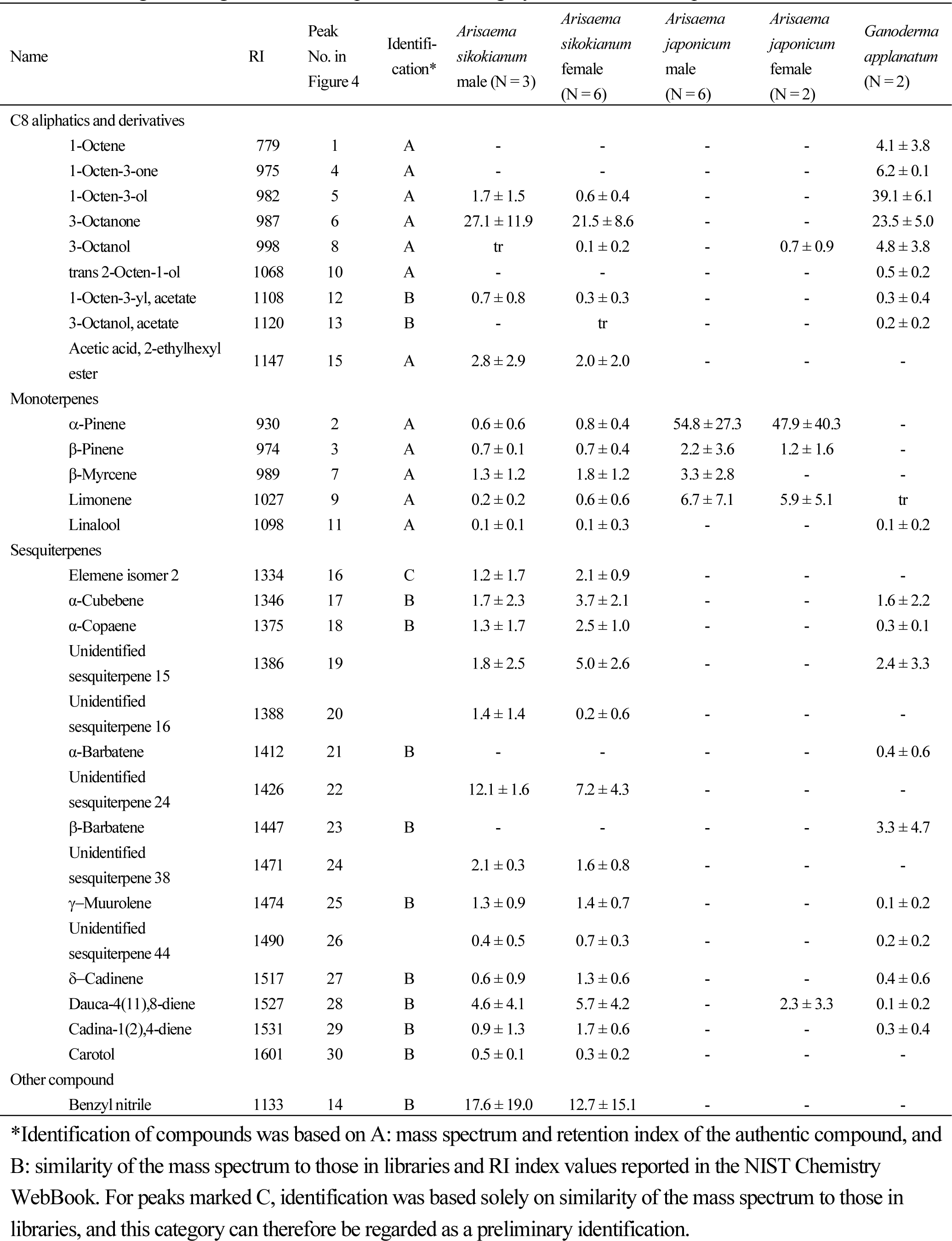
Representative volatile compounds detected in inflorescences of *Arisaema sikokianum* and *A. japonica* and fruiting bodies of *Ganoderma applanatum*. Percentages of the total ion chromatogram peak area are shown, from data averaged among individual samples for each category. The full dataset is provided in Table S6.

**Figure 4.**
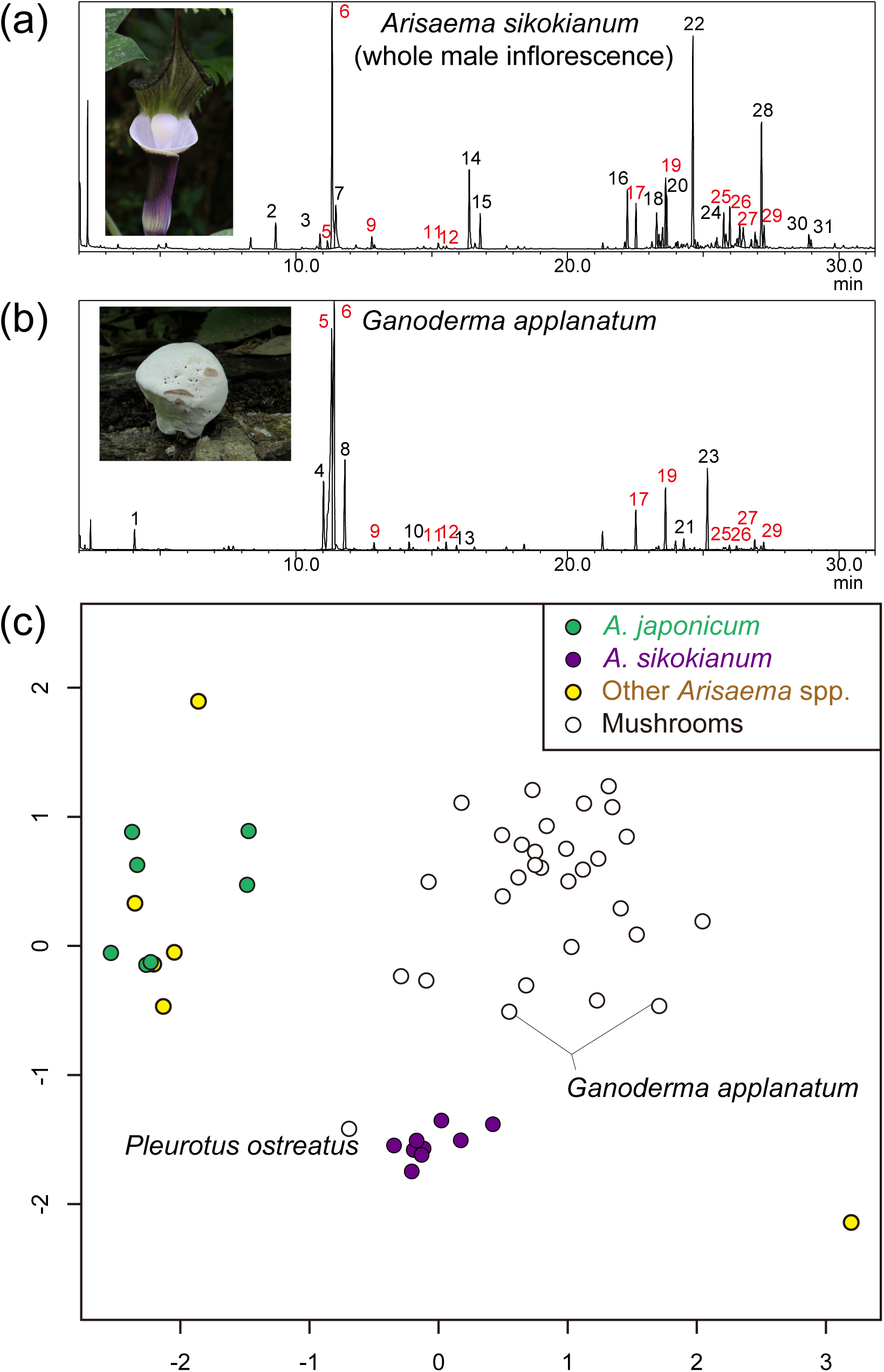
The volatile compositions of *Arisaema* spp. and the various mushrooms. (a and b) The total ion chromatogram of an inflorescence of *A. sikokianum* (a) and a fruiting body of *Ganoderma applanatum* (b). Only the principal compounds are shown with peak numbers, and the compounds in common between the two samples are shown in red. The peak numbers correspond to those shown in Table 3. (c) The nonmetric multidimensional ordination plot of the volatile compositions from samples of *Arisaema* spp. and various mushrooms. One Arisaema sample (*A. angustatum* 166181) was not included in the ordination analysis because it is extremely dissimilar to the other samples. *A. sikokianum, A. japonicum*, the other *Arisaema* spp., and the mushrooms are colored purple, green, yellow, and white, respectively.

We detected a total of 21 volatile compounds (5–13 compounds per sample) from eight *A. japonicum* individuals, 12 of which were monoterpenes (Table S6). Only *α*-pinene and limonene were common among the samples. The C8 aliphatics were rarely detected among the volatiles from *A. japonicum*, with the exception of two plants that emitted very small amounts of either 1,3-octadiene or 3-octanol. We further detected a total of 26 volatile compounds (0–17 compounds per sample) from 7 additional *Arisaema* species that are potentially sympatric with *A. sikokianum*, 15 of which were monoterpenes and none of which were C8 aliphatics (Table S6).

To examine whether *Arisaema* inflorescences chemically mimic mushrooms, we sampled volatiles from the headspace of 27 species of mushrooms and detected a total of 147 volatile compounds (4–43 compounds per sample; Table S6). Again, the compounds consisted mainly of monoterpenes, sesquiterpenes and C8 aliphatics. Only 3-octanone was common among all samples, and other C8 aliphatics (e.g. 1-octene, 1-octene-3-ol, 1-octene-3-one and 3-octanol) were common among the majority of the samples. Meanwhile, monoterpenes and sesquiterpenes were more variable and rarely common among the samples. Among these terpenoids, only a sesquiterpene *β*-barbatene was shared among >30% of the samples (10 fungal species), although the compound was not detected in the *Arisaema* samples.

To quantify the similarity of the volatile compositions from the *Arisaema* inflorescences and fungal fruiting bodies, we performed a nonmetric multidimensional scaling analysis based on Bray–Curtis dissimilarity calculated from a relative volatile composition matrix for each sample. We found that the level of dissimilarity of volatile compositions between *A. sikokianum* inflorescences and mushrooms (0.54–0.97, 0.81 on average) was generally lower than that between the other *Arisaema* spp. and mushrooms (0.83–1.00, 0.99 on average) or between *A. sikokianum* and the other *Arisaema* spp. (0.91–1.00, 0.97 on average). Accordingly, the volatile profiles of *A. sikokianum* were plotted close to those of the mushrooms, whereas those of *A. japonicum* and the other *Arisaema* spp. were plotted distantly to both *A. sikokianum* and the mushrooms (Figure 4*c*).

## Discussion

### *Pollination via mushroom mimicry in* A. sikokianum

The present study was motivated by a finding of one of the authors (YO) during the management of the plant collection in Tsukuba Botanical Garden; namely, that the inflorescence of *A. sikokianum* has a strong characteristic fragrance reminiscent of mushrooms. The striking morphological similarity of *A. sikokianum* to certain mushrooms, such as the young fruiting body of the bracket fungus *Ganoderma applanatum* (Pers.) Pat., suggests that the mimicry is likely to involve both visual and olfactory signals, as pointed out in an earlier study (Vogel and Martens 2000).

As expected, the volatile compositions of *A. sikokianum* inflorescences chemically resembled those of mushrooms (Figure 4*a* and *b*). First, they were dominated by C8 aliphatics common among the fruiting bodies of most of the fungal species examined, such as 3-octanone, 1-octene-3-ol and 1,3-octadiene. Moreover, they contained some monoterpenes and sesquiterpenes in common with wood-decaying fungi, such as *Ganorderma applanatum* (Tables 3 and S6), suggesting that the floral mimicry in *A. sikokianum* targets specific model fungal groups (e.g., some polypore fungi) rather than mushrooms in general.

The existence of mushroom mimicry in *A. sikokianum* was further supported by the striking similarity between the assemblages of the floral visitors and insect communities attracted to some mushroom species (Figure 3*b*), both of which were dominated by obligately mycophagous drosophilids (both males and females of *Mycodrosophila gratiosa* and other *Mycodrosophila* species). Judging from their high visitation frequency of both male and female inflorescences and ideal body size for effective pollination (see Materials and Methods), it is reasonable to identify *Mycodrosophila* spp. as the principal pollinator of *A. sikokianum*. We further confirmed that mushroom-like volatiles were emitted not only from the appendix but also from the spathe (Figure S1), and removal of the mushroom-like appendix in the *A. sikokianum* inflorescence resulted in only a partial decrease in pollinator visits without altering pollinator composition, which was dominated by *Mycodrosophila.* Note that the effect of appendix removal was not visible in 2017, which had greater variance in data compared to 2019, likely because the experiment was conducted later in the flowering season in 2017. Additionally, because we left the inflorescences of the control plants intact, our experimental design did not control for the wounding effect of appendix ablation. Therefore, the mushroom mimicry apparatus in *A. sikokianum* is likely to be achieved by both the mushroom-like appendix and the purple and white spathe.

Demonstrating pollination by floral mimicry is often challenging, since it requires an understanding of the pollinators’ perception without detailed knowledge of the life histories of hyperdiverse insects (especially dipterans). For example, stable artificial breeding of *Mycodrosophila* spp. has not yet been established, hindering behavioral or electrophysiological studies of the *A. sikokianum*–*Mycodrosophila* system. In this study, we demonstrated mushroom mimicry in *A. sikokianum* by comparing the pollinator assemblage of *A. sikokianum* to the insect communities attracted to various mushroom species in Japan. The latter data were available only because they were collected previously by one of the authors (Tuno 2001). Although data on mushroom-attracted insect communities were collected almost 20 years ago, their striking similarity to the pollinator assemblages of *A. sikokianum* supports the idea that a deceptive pollination system exploiting mushroom-attracted insect communities could be ecologically stable temporally and spatially. The present study therefore highlights the importance of quantitative data from local community-based studies regarding the insects attracted to specific substrates, such as mushrooms, carrion and animal feces, that are potentially mimicked by local plants.

### *Mushroom mimicry in* A. sikokianum *as a possible unique Batesian food-source mimicry*

The mushroom mimicry system in *A. sikokianum* is apparently most comparable with that reported in the neotropical orchids *Dracula* spp., which also exploit the mushroom-visiting behavior of mycophagous drosophilids (Policha et al. 2016, 2019). In *Dracula–*drosophilid systems, the pollinators are never physically trapped by the flowers and even receive some fitness benefits from the flowers, such as food, shelter, mating opportunities and, in rare cases, breeding; thus, the pollination system is somewhat mutualistic rather than purely deceptive (Policha et al. 2019). Conversely, in *A. sikokianum* the pollinators are unlikely to receive any fitness benefits because the plant species apparently lacks any floral reward, such as nectar, and even has a mechanism for imprisoning pollinators; pollination success is always coupled with this reproductive dead end for the pollinators. It is also unlikely that male inflorescences, from which the pollinators can escape, provide any fitness benefits to pollinators. This is because we seldom encountered pollinators inside the intact male inflorescences, despite their fairly frequent visits, indicating that the inflorescences do not provide any practical benefit that would retain pollinators. Therefore, the pollination system of *A. sikokianum* may be unique in that it is an instance of purely deceptive mushroom mimicry.

Pollination by mushroom mimicry has often been discussed under the context of oviposition-site mimicry (Johnson and Schiestl 2016), as some of the known examples of this pollination mode involve the oviposition behavior of the pollinators (Vogel 1978, Sugawara 1988). However, based on our observations of *Mycodrosophila* spp., the principal pollinator of *A. sikokianum*, did not lay eggs inside the spathe of *A. sikokianum. Mycodrosophila* spp. visit fruiting bodies of some bracket fungi, such as *Ganoderma applanatum*, only to feed on spores on the surface (Tuno 1999, Kobayashi et al. 2017), although fragmentary information suggests that they breed in the fruiting bodies of other fungi (Kimura et al. 1977). Therefore, *A. sikokianum* likely exploits adult foraging behavior rather than the oviposition of the pollinators, so it might be considered a special case of Batesian food-source mimicry.

### *Evolution of mushroom mimicry as a possible mode of ecological speciation in* Arisaema

We revealed that *A. sikokianum* has a pollinator assemblage distinct from other *Arisaema* spp., probably as a consequence of its unique visual and chemical mushroom mimicry traits. We are aware of at least 15 *Arisaema* spp. whose pollinators are known, but none are dominated by Drosophilidae pollinators or have typical mushroom mimicry traits (Vogel and Martens 2000, Nishizawa et al. 2005, Barriault et al. 2010, Tanaka et al. 2013, Kakishima and Okuyama 2018). We further confirmed that floral volatile profiles of the other *Arisaema* spp. that are potentially sympatric to *A. sikokianum*, including *A. japonicum*, bore no resemblance to those of mushrooms (Figure 4*c*). These results indicate that mushroom mimicry is unusual within this genus, although this view seems somewhat contradictory with traditional views on *Arisaema* pollination.

Mushroom mimicry has long been suggested as a pollination mechanism for this genus (Vogel and Martens 2000), since most reported pollinators for *Arisaema* spp., and for *A. japonicum* in the present study, are fungus gnats (Sciaridae and Mycetophilidae) (Sasakawa 1993, 1994a, b, Vogel and Martens 2000). Fungus gnats are often reported to be mycophagous in their larval stage (but see Freeman 1983 and Okuyama et al. 2018 for a discussion of herbivory among these insects). However, several studies have shown that adult fungus gnats often feed on nectar from flowers (Ackerman and Mesler 1979, Okuyama et al. 2004, 2008), suggesting that some volatile signals such as monoterpenes (e.g., linalool and its derivatives in *Mitella* spp.) (Okamoto et al. 2015) may attract these insects, as well as visual signals such as the greenish or reddish color (Mochizuki and Kawakita 2017) of fungus-gnat-visited flowers. By contrast, *Mycodrosophila* spp., the principal pollinator of *A. sikokianum*, never visit flowers for nectar, whereas both male and female flies visit various mushrooms for courtship semaphoring, food, shelter, territory defense and mating (Policha et al. 2019). Collectively, these observations imply that the mushroom mimicry in *A. sikokianum* is derived from Batesian food-source mimicry involving fungus, which might be prevalent in the genus *Arisaema*.

It is noteworthy that pollination via mushroom mimicry in *A. sikokianum* likely functions as an efficient reproductive barrier from other sympatric, closely-related *Arisaema* spp. such as *A. japonicum*, because there is little overlap between them in pollinator assemblages Matsumoto et al. (*in press*) also demonstrated strong pollinator isolation between *A. sikokianum* and *A. tosaense* in their sympatric populations. Accordingly, it is possible that the evolutionary acquisition of mushroom mimicry traits in the *A. sikokianum* lineage coincided with ecological speciation, as most species of genus *Arisaema* section *Pistillata*, including *A. sikokianum*, are cross-compatible based on extensive artificial cross experiments (Murata et al. 2018). Although the species most closely related to *A. sikokianum* is currently unknown, it is clear that the speciation event took place very recently, since a recent phylogenetic study of the genus using ∽2.8 kb chloroplast DNA found little genetic variation among 31 spp., including *A. sikokianum*, belonging to the same section (*Pistillata*) (Ohi-Toma et al. 2016). Future comparative studies between *A. sikokianum* and its sister species in the fields of ecology, genetics or physiology may provide an excellent opportunity to study the mechanisms of coordinated acquisition of elaborate mushroom mimicry traits, and to test the hypothesis of ecological speciation associated with mushroom mimicry, which is one of the most bizarre phenomena in floral evolution.

## Acknowledgments

The authors thank Masanori Toda for the identification of drosophilids, Hiroko Murata for the living material of *Arisaema yamatense*, and Chiemi Takaboshi and Yumiko Suzuki for supporting the DNA barcoding of pollinators. We also thank the anonymous reviewers for their helpful comments on an earlier draft of the manuscript. This work was supported by JSPS KAKENHI Grant Numbers 15H05604 and 19H03292, and the 28th Botanical Research Grant of the ICHIMURA Foundation For New Technology (to Y.O.). We certify that the present research was conducted in conformance with all applicable laws regarding the endangered species (*A. sikokianum*).

---

**Figure S1.**
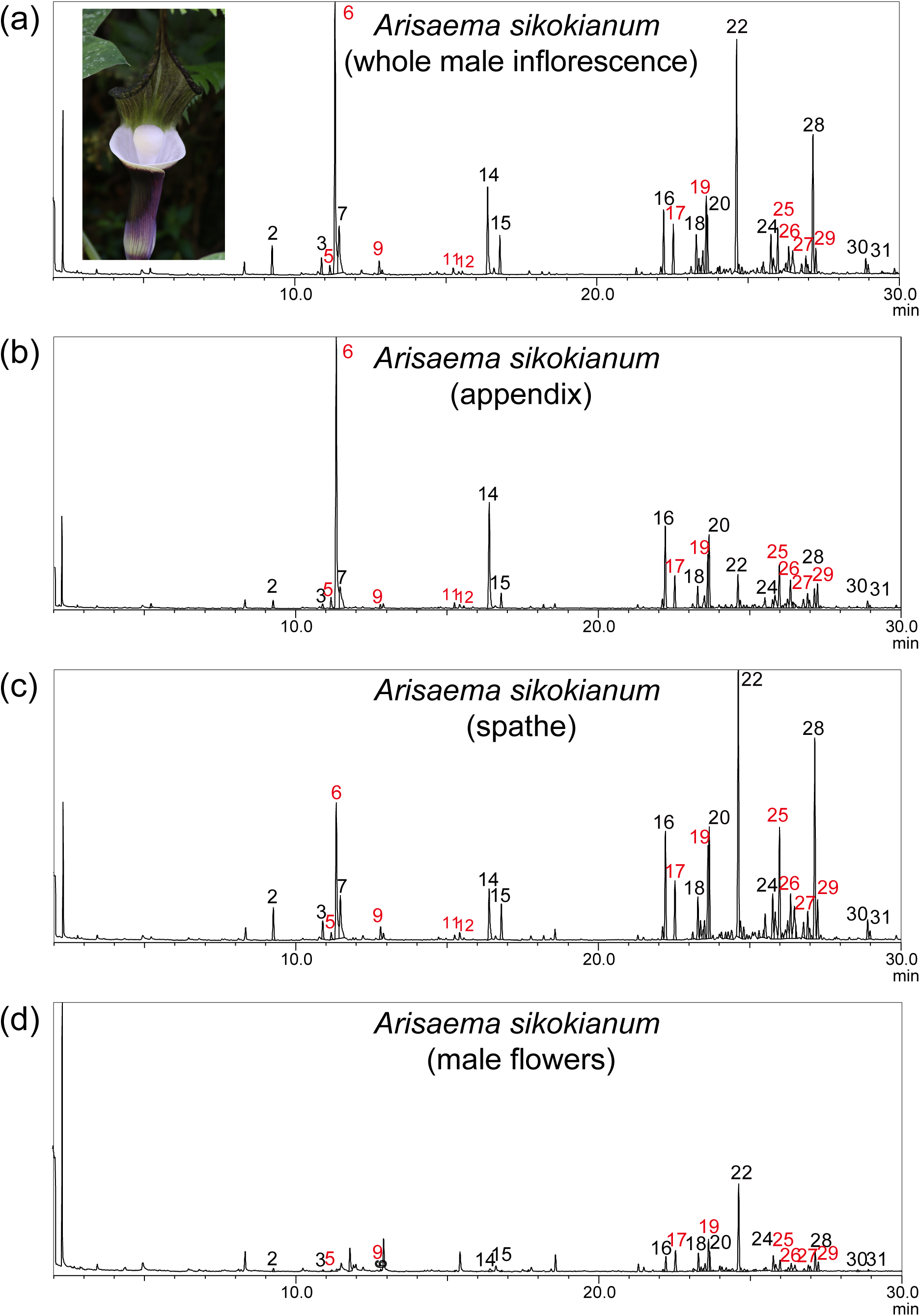
The total ion chromatogram of an inflorescence of *A. sikokianum* analyzed separately in different parts. (a) Whole inflorescence (the same chromatogram as in Fig. 4. a). (b) Appendix. (c) Spathe. (d) Male flower cluster.

